# Bacterial competition systems share a domain required for inner membrane transport of the nuclease bacteriocin pyocin G

**DOI:** 10.1101/2021.11.17.469063

**Authors:** Iva Atanaskovic, Connor Sharp, Cara Press, Renata Kaminska, Colin Kleanthous

**Author notes:** **Author Contributions:** I.A., C.S., C.P. and R.K. performed research. I.A. and C.S. analyzed data. I.A., C.S. and C.K. designed research. I.A., C.S. and CK. wrote the paper. C.K. conceived the project. **Competing Interest Statement:** The authors have declared that no competing interests exist.

## Abstract

Bacteria exploit a variety of attack strategies to gain dominance within ecological niches. Prominent amongst these are contact-dependent inhibition (CDI), type VI secretion (T6SS) and bacteriocins. The cytotoxic endpoint of these systems is often the delivery of a nuclease to the cytosol. How such nucleases translocate across the cytoplasmic membrane of Gram-negative bacteria is unknown. Here, we identify a small, conserved, 15-kDa domain, which we refer to as the inner membrane translocation (IMT) domain that is common to T6SS and bacteriocins and linked to nuclease effector domains. Through fluorescence microscopy assays using intact and spheroplasted cells, we demonstrate that the IMT domain of the *Pseudomonas aeruginosa* specific bacteriocin pyocin G (PyoG) is required for import of the toxin nuclease domain to the cytoplasm. We also show that translocation of PyoG into the cytosol is dependent on inner membrane proteins FtsH, a AAA+ATPase/protease, and TonB1, the latter more typically associated with transport of bacteriocins across the outer membrane. Our study reveals that the IMT domain directs the cytotoxic nuclease of PyoG to cross the cytoplasmic membrane and, more broadly, has been adapted for the transport of other toxic nucleases delivered into Gram-negative bacteria by both contact dependent- and contact-independent means.

**Importance:** Nuclease bacteriocins are potential antimicrobials for the treatment of antibiotic resistant bacterial infections. While the mechanism of outer membrane translocation is beginning to be understood, the mechanism of inner membrane transport is not known. This study uses PyoG as a model nuclease bacteriocin and defines a conserved domain which is essential for inner membrane translocation and which is widespread in other bacterial competition systems. Additionally, the presented data links two membrane proteins, FtsH and TonB1, with inner membrane translocation of PyoG. These findings point to the general importance of this domain to the cellular uptake mechanisms of nucleases delivered by otherwise diverse and distinct bacterial competition systems. The work is also of importance for the design of new protein antibiotics.

## Introduction

Bacteria deploy various contact dependent and independent competition systems to compete for space and resources (1). These systems deliver toxic effectors to the cell envelope or the cytoplasm of bacterial competitors. While the mechanism of effector delivery varies between different competition systems, effector structures and killing mechanisms can be conserved (2). Often, competition system effectors are nucleases that get transported across the cell envelope to degrade cytoplasmic nucleic acids. These folded proteins cross the multi-layered cell envelope via poorly understood translocation mechanisms.

Bacteriocins of Gram-negative bacteria are protein antibiotics deployed as weapons for contact-independent bacterial competition. Bacteriocins have a potential to be developed into antibiotics, for the treatment of infections resistant to conventional antimicrobials (3). Bacteriocins can kill bacterial cells via different mechanisms: by forming pores in the inner membrane; by inhibiting peptidoglycan biosynthesis through lipid II degradation; by DNA or RNA degradation. Bacteriocin producing strains are immune to their own bacteriocins due to the production of immunity proteins that inhibit the bacteriocin’s killing activity. Sensitive strains do not produce the immunity protein or produce an immunity protein specific for a different bacteriocin, but nevertheless have the appropriate translocation machinery through the cell envelope. In the case of nuclease bacteriocins, which penetrate the cytoplasm to degrade nucleic acids, this translocation machinery spans both the outer and the inner membranes (4).

Pyocins are bacteriocins that target and kill *Pseudomonas aeruginosa*. Pyocin G (PyoG) is a nuclease pyocin active against *P. aeruginosa* clinical isolates (5). It is composed of an unstructured N-terminus followed by a receptor binding domain (5), a conserved middle domain which is essential for its killing activity (6), and a cytotoxic nuclease domain (7). The receptor binding domain is required for outer membrane translocation of PyoG (5). The role of the conserved central domain is unknown, but the fact that it is absent from bacteriocins with periplasmic targets and present in bacteriocins that cleave nucleic acids (Pfam domain PF06958.7) suggests that it may have a role in inner membrane transport (8). Here, we show that this domain is required for PyoG translocation to the cytosol.

PyoG enters the periplasm by binding and translocating through Hur in the outer membrane (5). Hur is a 22-stranded β-barrel TonB dependent transporter (TBDT) that ordinarily transports hemin into the cell following engagement with TonB1 in the inner membrane, in conjunction with the stator complex ExbB-ExbD and the proton motive force (PMF). TonB1 activates hemin transport following association with Hur’s periplasmically-located TonB box epitope. PyoG parasitizes hemin uptake and in so doing delivers its own TonB box to the periplasm which engages with TonB1, allowing the toxin to be pulled into periplasm.

TonB-dependent outer membrane translocation of pyocins is well-understood (5, 9, 10). By contrast, how these toxins translocate from the periplasm to the cytoplasm is unknown. In the case of nuclease colicins, which are *E. coli*- specific bacteriocins, inner membrane transport requires both the ATPase and protease functions of FtsH (11). FtsH is a hexameric AAA+ ATPase/protease in the inner membrane, where it is involved in protein quality control (12). FtsH proteolytically processes nuclease colicins as they pass into the cytoplasm (13). More recently, FtsH has also been linked to the killing activity of PyoG (5) demonstrating a broader involvement in bacteriocin uptake.

In the present work, we address the poorly understood area of bacteriocin transport across the inner membrane. Using a newly developed import assay, whereby the uptake of fluorescently labelled PyoG into *P. aeruginosa* spheroplasts is monitored, we map the requirements for pyocin transport to the cytoplasm. Using this assay, we demonstrate that a small domain found in all nuclease bacteriocins as well as other bacterial competition systems is absolutely required for transport.

## Results and Discussion

### Inner membrane translocation of nucleases in bacterial competition systems is associated with a ubiquitous domain

Using bioinformatics, we have previously identified a highly conserved bacteriocin domain that is absent from pore forming bacteriocins but associated exclusively with competition systems that transport nucleases to the cytoplasm (8). We term this domain the Inner Membrane Translocation (IMT) domain. The IMT domain is annotated in the PFAM database as the pyocin-S domain (PF06958.7). The main structural features of the domain are two anti-parallel β-sheets that give the domain an L-shape (Figure 1 - A). Using a more extensive informatics search we found the IMT domain across several orders of Gammaproteobacteria (Figure 1 - B). Surprisingly, apart from being conserved amongst nuclease bacteriocins, this domain is also found in Type 6 Secretion System (T6SS) effectors (Figure 1 - C). These toxins, like nuclease bacteriocins, translocate across the inner membrane to kill competitors. Unlike bacteriocins however, which are diffusible toxins, T6SS effectors are contact dependent and delivered by a contractile needle that punctures the outer membrane. We found several characteristic PFAM domains of these toxin systems co-occur with the IMT domain (Figure 1 - D) – the HNH nuclease domain found in DNase bacteriocins and T6SS effectors, and the rRNase and tRNase bacteriocin domains. We conclude that the IMT domain is found in competition system effectors that express their cytotoxic activity in the cytoplasm of target cells regardless of the means by which the toxin initially penetrates the outer defenses of the Gram-negative bacterium.

**Figure 1.**
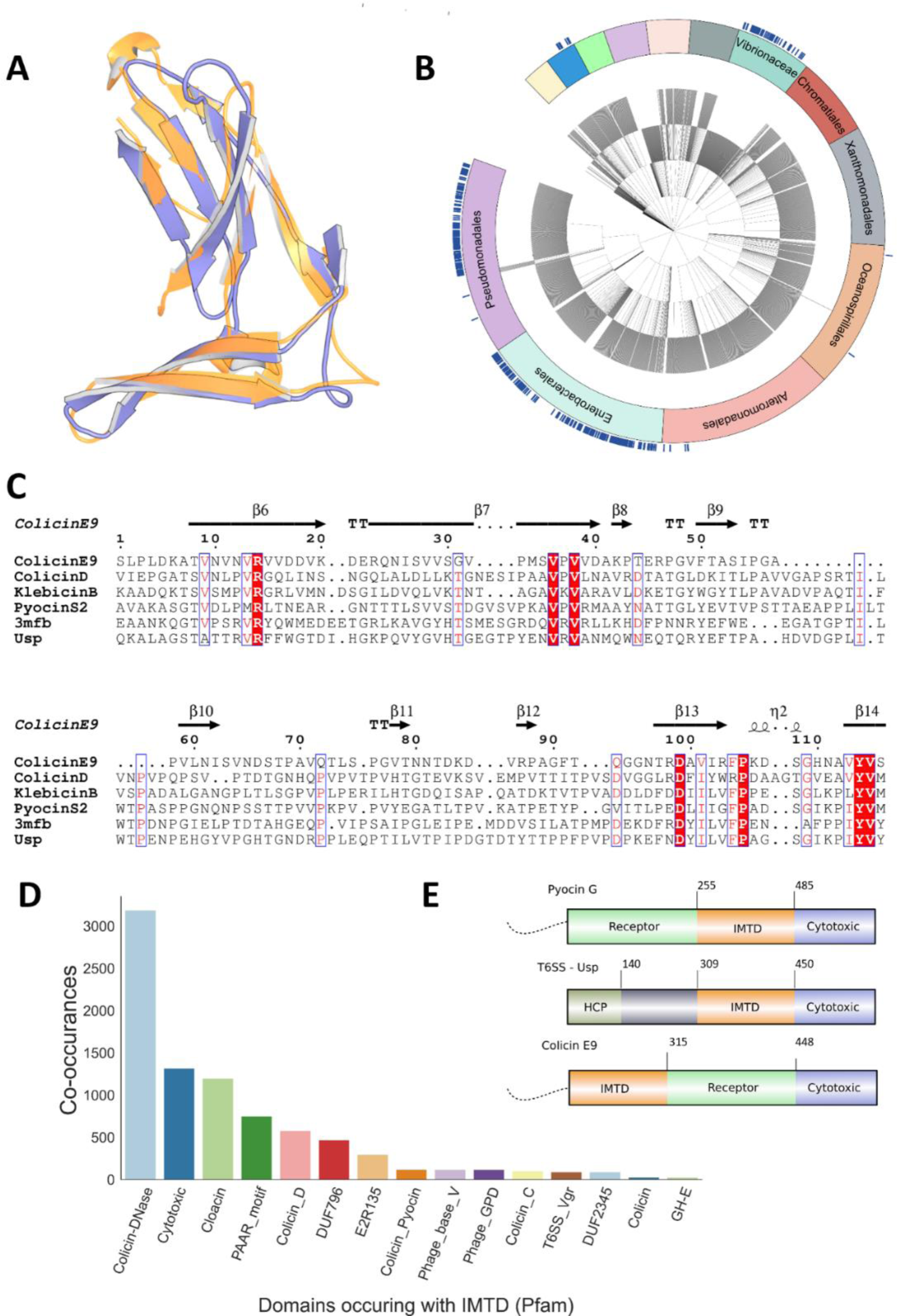
The IMT domain is a conserved structural domain found in multiple orders within Gammaproteobacteria and associated with multiple toxin systems. **A** - Structure of the conserved beta-sheets of the IMT domain from colicin D (blue - pdb: 5ZNM) and a T6SS effector (gold - pdb: 3MFB). **B** - Taxonomy of Gammaproteobateria within uniref100. Blue ticks indicate species which contain at least 1 IMT domain containing protein. IMT domain proteins are prevalent in Gammaproteobacteria though restricted to certain orders (Unlabelled taxa are: Cellvibrionales – Grey, Thiotrichales- Peach, Pasteurellaceae - Purple, Legionellales – Green, Aeromonadales – Blue, Methylococcales – Yellow). **C** - Alignment of the IMT domain domain from bacterial toxins including colicins and other nuclease bacteriocins and T6SS effectors (3MFB and Usp – Uropathic Specific Protein). **D** - Pfam domains which co-occur in proteins with IMT domain. IMT domain is found in proteins which contain numerous toxin effectors or structural domains. (Colicin-DNase – common HNH nuclease found in DNase bacteriocins and T6SS effectors, Cytotoxic – rRNase toxin found in bacteriocins and T6SS effectors, Colicin_D – tRNase found in bacteriocins and T6SS effectors, DUF796 (T6SS_HCP) – structural motif found in T6SS effectors, E2R135 – receptor domain for certain colicins, Colicin_Pyocin – a domain found in the immunity proteins for many DNase bacteriocins. Phage_base_V/Phage_GPD – phage/T6SS structural proteins, Colicin_C – tRNase found in bacteriocins and T6SS effectors, T6SS_Vgr – structural domain of the T6SS, DUF2345 – domain of unknown function associated with the T6SS, Colicin – pore forming domain found in many colicins, all proteins identified with this domain had similarity to Colicin B, GH-E – HNH family of nucleases). **E** – The domain organisation of PyoG (5). For comparison, the position of the IMT domain is shown for a T6SS nuclease effector (Usp) and a nuclease colicin (ColE9).

### Inner membrane translocation of PyoG requires the IMT domain

Having established that the IMT domain is associated with inner membrane translocation of toxin nucleases in bacteria, we sought to test its requirements for import of PyoG into *P. aeruginosa*. PyoG is a 640 amino acid toxin comprised of an N-terminal receptor binding domain that engages Hur and enables transport to the periplasm and a C-terminal nuclease domain that elicits cell death (5). Between them is the IMT domain which comprises residues 256-485 (Figure 1 – E). We developed a fluorescence microscopy assay to dissect the involvement of the IMT domain in transport. Pyocins can be readily conjugated with fluorescent dyes and used for labelling live *P. aeruginosa* cells that express components of the pyocin translocation machinery in microscopy experiments. Trypsin protection is then used to distinguish imported from surface bound molecules(9, 10). We generated different PyoG constructs, each containing a unique cysteine at the C-terminus of the construct that was conjugated to AlexaFluor (AF) 488 and assessed the ability of each construct to be imported in intact cells and spheroplasts (Figure 2 – A). Spheroplasts were generated by lysozyme/EDTA treatment that permeabilises the outer membrane and peptidoglycan layer (14, 15, 16). After labelling with fluorescent pyocin, cells were exposed to trypsin to remove any surface exposed pyocin that was not translocated across the cell envelope. In intact cells, the outer membrane is not permeable to trypsin(9, 10). Therefore, in intact cells translocation across the outer membrane is sufficient to protect a pyocin construct from degradation with trypsin. In spheroplasts, the periplasm is exposed to trypsin (14), and the pyocin construct must translocate across the inner membrane to be protected from degradation by trypsin. Residual fluorescence after trypsin treatment is therefore an indication of outer membrane, and potentially inner membrane transport, in intact cells. In the case of spheroplasts, it is an indication of inner membrane transport. To test if nontranslocated pyocin gets degraded by trypsin under the conditions of our experiments, a pyocin S2-GFP chimera was used (Supplementary Figure 1 - A). This construct is comprised of the pyocin S2 N-terminal domain (residues 1-209) translationally fused to GFP. GFP acts as a plug, blocking the import of the pyocin (10). Trypsin treatment removed the PyoS2- GFP fluorescent signal entirely in both intact cells and spheroplasts (Supplementary Figure 1 – A, B). A further control was undertaken to ascertain if the outer membrane translocation step can be bypassed using spheroplasts. *Δhur*, a PAO1 transposon mutant that lacks the PyoG receptor (5), was exposed to fluorescent PyoG. Intact cells of this mutant were not labelled with the pyocin, since Hur is essential for binding of PyoG to the surface of *P. aeruginosa* cells (5). Generation of spheroplasts was sufficient to allow *Δhur* labelling with fluorescent PyoG. Additionally, PyoG was protected from degradation by trypsin in *Δhur* spheroplasts (Supplementary Figure 1 – D, E). Therefore, the Hur-dependent outer membrane translocation step can be bypassed under these tested experimental conditions, confirming that *P. aeruginosa* spheroplasts could be used to study inner membrane translocation.

**Figure 2.**
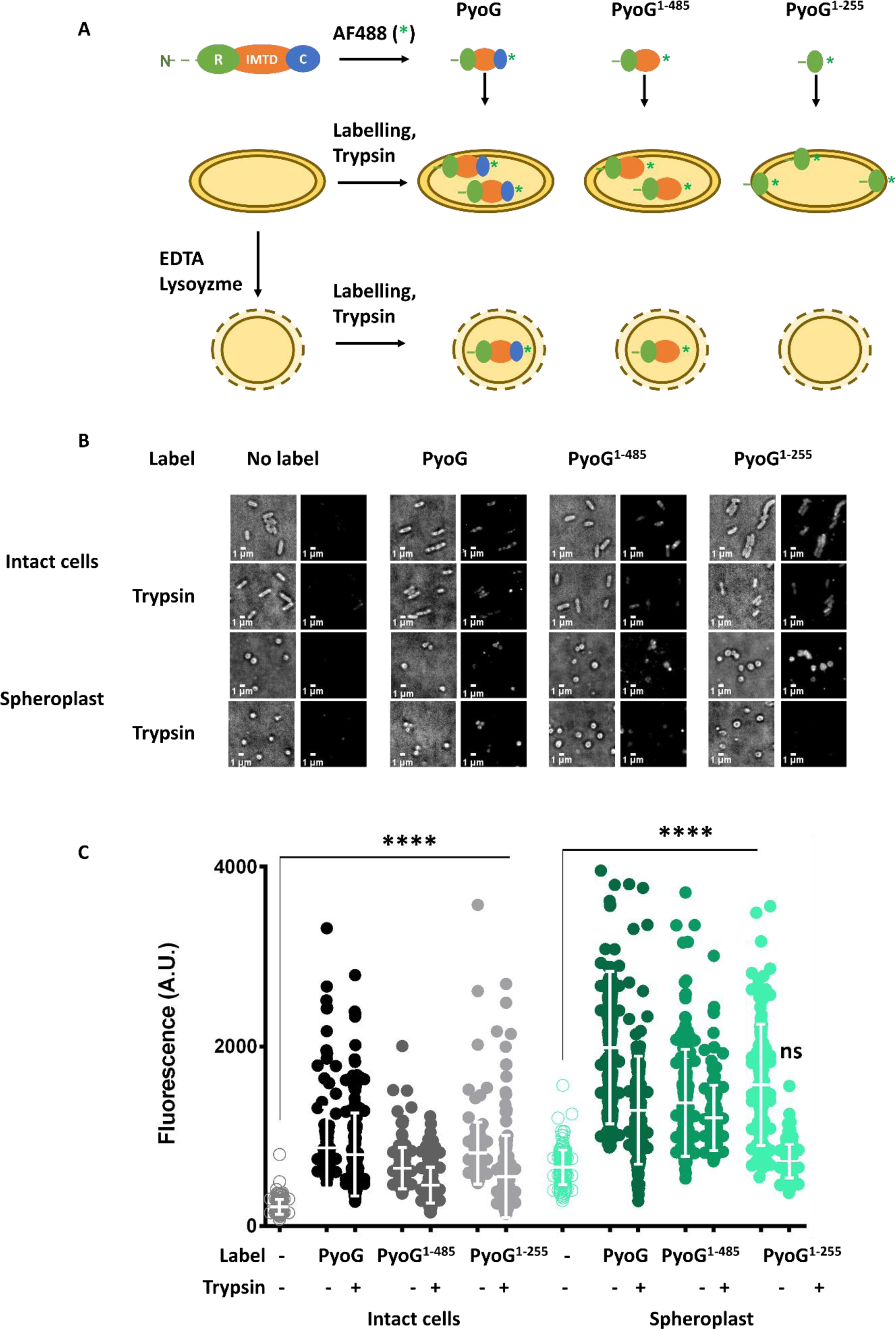
Import and localisation of fluorescent PyoG constructs. **A** – a fluorescence microscopy experiment set-up, used for localisation of PyoG constructs in *P. aeruginosa* cells. All constructs are conjugated to AF488 (represented with a green *) via a C-terminal cysteine. Intact cells, or spheroplasts generated by lysozyme/EDTA treatment, are exposed to 2 µM fluorescent PyoG. Unbound and untranslocated pyocin constructs are removed by trypsin treatment. Residual fluorescence in intact cells is an indication of outer membrane, and possibly inner membrane translocation. Since the spheroplast outer membrane and periplasm are disrupted by EDTA and lysozyme, the periplasm is exposed to trypsin(14, 16). Therefore, residual fluorescence in spheroplasts is an indication of inner membrane translocation. **B** - Representative micrographs before and after trypsin treatment are shown. All snapshots were adjusted to the same contrast value. Tested constructs are – full length PyoG, PyoG^1-485^ lacking the cytotoxic domain, and PyoG^1-255^ lacking the cytotoxic domain and the IMT domain. All constructs are trypsin protected in intact cells, indicating outer membrane translocation. In spheroplasts, full length PyoG and PyoG^1-485^ are trypsin protected, but PyoG^1-255^ is not. Therefore, the IMT domain is required for inner membrane translocation of PyoG. **C** - Average fluorescence intensities were measured for 150 cells per condition. Mean of three biological replicates with standard deviations are shown. Fluorescence intensities for labelled and trypsin treated groups in each condition are compared to the unlabelled control. **** represents a P value below 0.0001 in the Kruskal-Wallis Test, and ns represents no significant difference or lack of fluorescent labelling.

The cellular localisation of the following PyoG constructs were tested: full length PyoG, PyoG^1-485^ which lacks the cytotoxic domain, and PyoG^1-255^ which lacks the cytotoxic and IMT domains (Figure 2 – A). Constructs were expressed and purified as previously described (5). Their fold and stability were tested by circular dichroism (Supplementary Figure 1 – A) and differential scanning calorimetry (Supplementary Figure 1 - B); all appeared folded at room temperature. All three constructs contained the receptor (Hur-) binding domain of PyoG (residues 1-255) and were protected from degradation by trypsin in intact cells (Figure 1 – B, C), which indicated that these constructs translocated across the outer membrane. This is in accordance with previously published work, where a pyocin receptor binding domain was sufficient to protect the pyocin from degradation by extracellular trypsin added to live *P. aeruginosa* cells(9, 10). However, only full length PyoG and PyoG^1-485^ were trypsin protected in spheroplasts (Figure 1 – B, C), indicating that these two constructs were capable of translocating across the inner membrane. Trypsin protection of PyoG^1-485^ also implied that the cytotoxic domain of PyoG was not essential for inner membrane translocation, which has previously been suggested to be a requirement for colicin transport (11). PyoG^1-255^, which lacks the IMT domain, was not trypsin protected in *P. aeruginosa* spheroplasts. We therefore conclude that the IMT domain is required for PyoG to cross the inner membrane.

### Inner membrane translocation of PyoG requires FtsH and TonB1

PyoG killing activity has previously been linked to Hur, the outer membrane receptor, TonB1, the inner membrane protein that provides energy for outer membrane translocation, and FtsH in the inner membrane (5). Since FtsH is not essential in *P. aeruginosa*, we used PAO1 as a model organism to test if FtsH is required for outer and/or inner membrane translocation of PyoG. PAO1 wt and *ΔftsH* cells were exposed to fluorescent PyoG^1-485^ and to trypsin, as described in Figure 2. PyoG^1- 485^ was trypsin protected in *ΔftsH* intact cells (Figure 3 – A, B), demonstrating that outer membrane translocation of PyoG does not require FtsH. On the other hand, PyoG^1-485^ was not trypsin protected when added to *ΔftsH* spheroplasts (Figure 3 – A, B), indicating that FtsH was required for inner membrane translocation of PyoG. Additionally, when the *ftsH* deletion was complemented from a plasmid, PyoG killing activity (Supplementary Figure 3 - A), and inner membrane translocation (Figure 3 – A, B) were restored. Complementation with FtsH H416Y, a mutant of FtsH that does not have protease activity (17), did not restore PyoG killing activity (Supplementary Figure 3 - A), or inner membrane translocation (Figure 3 – A, B). Therefore, proteolytically active FtsH is required for PyoG to cross the inner membrane.

**Figure 3.**
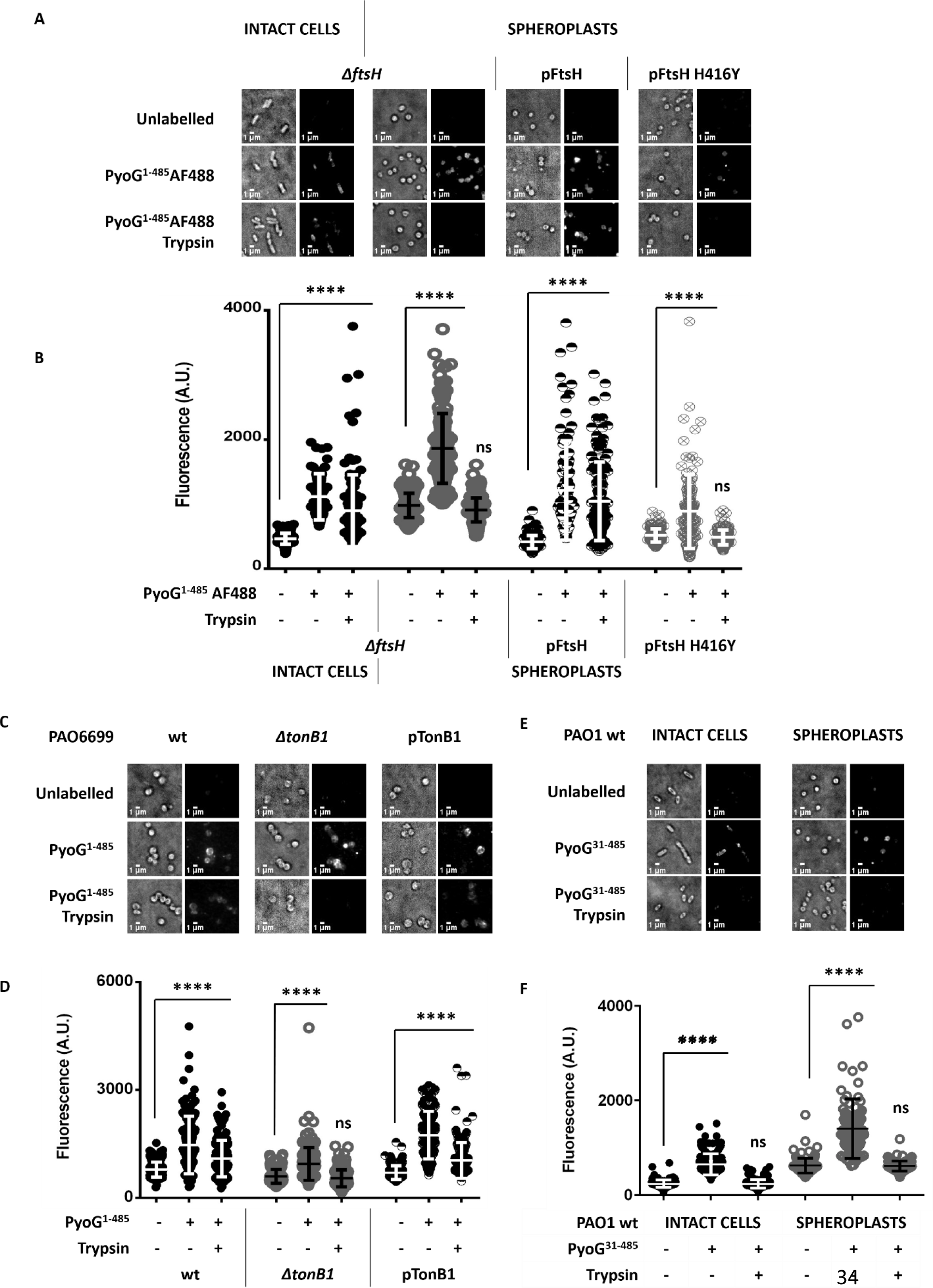
Inner membrane translocation of PyoG requires FtsH and binding to TonB1. PyoG^1-485^ is conjugated to AF488 via a C-terminal cysteine and added to *P. aeruginosa* spheroplasts at 2 µM. Nontranslocated pyocin is removed by trypsin treatment. **A, B** - FtsH is required for inner membrane translocation of PyoG. PyoG is trypsin protected in intact *ΔftsH* cells, but not in *ΔftsH* spheroplasts. This phenotype is complemented with FtsH expressed from a plasmid (pFtsH). No PyoG translocation is measured if the protease activity of FtsH has been disrupted with a point mutation (FtsH H416Y). Therefore, proteolytically active FtsH is specifically required for the inner membrane translocation of PyoG. **C, D** - Labelling of *ΔtonB1* spheroplasts with fluorescent PyoG. No residual labelling is measured after trypsin treatment, which indicates that TonB1 is required for inner membrane translocation of PyoG. PyoG import into spheroplasts is restored if TonB1 is expressed from a plasmid (pTonB1). **E, F** – The TonB binding box of PyoG, located in the unstructured N-terminus, is required for PyoG import. PyoG^31-485^ lacks the first 30 residues of PyoG and is conjugated to AF488 via a cysteine in the C- terminus. This deletion eradicates the killing activity of PyoG and its binding to the periplasmic domain of TonB1 (Supplementary Figure 4). This deletion also disrupts both the outer and the inner membrane import of PyoG, since the PyoG^31-485^ label is not trypsin protected in both intact cells and *P. aeruginosa* spheroplasts. Representative micrographs before and after trypsin treatment are shown (**A, C, E, G**). All snapshots were adjusted to the same intensity scale. Average fluorescence intensities were measured for 150 cells per condition (**B, D, F, H**). Mean of three biological replicates with standard deviations are shown. Fluorescence intensities for labelled and trypsin treated groups in each condition are compared to the unlabeled control. **** represents a P value below 0.0001 in the Kruskal-Wallis Test, and ns represents no significant difference or lack of fluorescent labelling.

We also tested if TonB1, a protein previously linked to pyocin outer membrane translocation(9, 10), is required for PyoG import into spheroplasts. This protein was required for inner membrane translocation of PyoG, since PyoG^1-485^ was not trypsin protected in *ΔtonB1* spheroplasts (Figure 3 – C, D). Complementation of *ΔtonB1* with TonB1 expressed from a plasmid restored PyoG killing activity (Supplementary Figure 3 - B) and inner membrane import (Figure 3– C, D). We also identified a region of PyoG that is involved in binding to TonB1 (Supplementary Figure 4) and demonstrated that this region is required for both outer and inner membrane import (Figure 3 – E, F). Pyocins S5 (9) and S2 (10) bind TonB1 via a TonB box located in the unstructured N-terminus. Therefore, we deleted the first 30 residues in the unstructured N-terminus of PyoG to test if this deletion affects TonB1 binding. The deletion, which renders PyoG inactive against *P. aeruginosa* (Supplementary Figure 4 – C), disrupted binding of PyoG to TonB1 (Supplementary Figure 4 – A), but it did not affect binding to the receptor Hur (Supplementary Figure 4 – B), confirming that the TonB box of PyoG is in the unstructured N-terminus. Unlike PyoG^1-485^, PyoG^31-485^ did not translocate into intact cells or spheroplasts (Figure 3 – E, F), demonstrating that the TonB box is required for both the outer and the inner membrane translocation steps.

### Model of nuclease bacteriocin transport across the inner membrane

The present study shows that the IMT domain, TonB1, and the ATPase/protease FtsH, are all essential for the inner membrane translocation step of PyoG. Taken together with previous studies that demonstrate FtsH dependent processing of nuclease colicins during import(18, 13), we propose a model of PyoG inner membrane translocation (Figure 4). The PyoG receptor binding domain (residues 1-255) binds to Hur, that interacts with TonB1 via its plug domain. The TonB box in the bacteriocin N-terminus also associates with TonB1, likely pulling through the TBDT to enter the periplasm in its entirety (Figure 5 - A; 10). PyoG associates with TonB1 via its TonB box but whether this interaction also exploits the PMF-dependence of TonB1, or other interaction partners is currently unknown. One possible role of the TonB1 interaction may be to localise PyoG close to the inner membrane from where the IMT domain either interacts directly with the membrane or even with FtsH itself for transfer across the membrane and proteolytic processing(18, 13). However, we were unable to demonstrate a direct interaction between FtsH and PyoG by pull-down assay (data not shown). It is also uncertain whether the entire IMT domain region of PyoG gets imported into the cytoplasm. Previous studies on colicins suggest that colicin D, E3 (18), E2, and E7 (13) undergo proteolytic processing during import. A cleavage site positioned within the IMT domain was defined for these colicins. Other studies on colicins have suggested that direct interaction of the transported nuclease with the cytoplasmic membrane is a requirement for transport to the cytoplasm(19, 20). However, PyoG constructs lacking the nuclease are still able to translocate (Figure 2). The IMT domain was sufficient to initiate inner membrane translocation, and potentially this domain also inserts into the inner membrane. Additionally, it is possible that the IMT domain initiates inner membrane transport through a mechanism involving other inner membrane proteins. The IMT domain of the nuclease colicin, ColD, has previously been shown to interact with inner membrane proteins essential for the ColD killing activity. The IMT domain of colicin D, which shares 26 % sequence identity with that of PyoG, has been shown to bind TonB(21, 22) as well as the signal peptidase, LepB in the inner membrane. Therefore, it is possible that the PyoG IMT domain also has additional interaction partners in the inner membrane.

**Figure 4.**
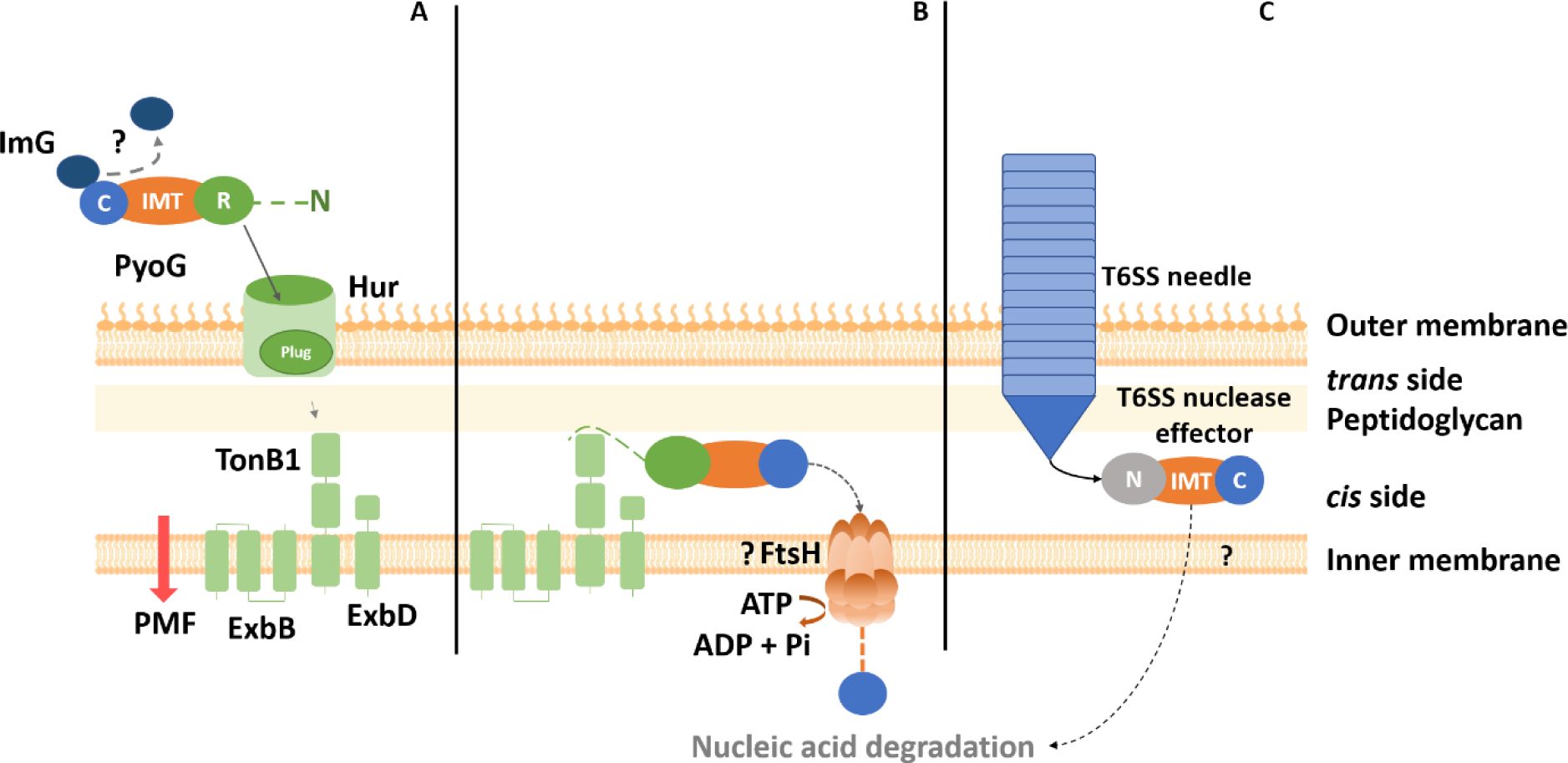
Probable import mechanism of nuclease pyocin and T6SS effector. **A** – PyoG uses Hur, a TBDT, as its outer membrane receptor and translocator (5). The pyocin is composed of an unstructured N terminus (dashed line), a receptor binding domain (R), a conserved inner membrane translocation domain (IMT domain), and a cytotoxic nuclease domain (C). Like in the case of nuclease colicins (28), the immunity protein (Im) probably disassociates from the C domain during translocation. The plug domain of Hur interacts with the periplasmic domain of TonB1 (5). The binding of TonB1 and the pyocin R domain to the TBDT induces a conformational change that dislocates the plug of the TBDT, allowing the passage of the pyocin through the TBDT (10). The TonB box, located in the N-terminus of PyoG, is essential for this translocation step, probably because TonB1 pulls PyoG through Hur and into the periplasm(9, 10). **B** – Translocation of PyoG from the periplasm into the cytoplasm also requires TonB1 binding. The exact role of TonB1 in inner membrane transport remains unknown. Potentially, TonB1 positions the pyocin on the surface of the inner membrane so it can interact with other proteins involved in inner membrane transport. This translocation step requires the IMT domain, a conserved domain present in all nuclease bacteriocins. Inner membrane transport of PyoG requires FtsH, an inner membrane ATPase/protease previously associated with the killing activity of PyoG (5) and nuclease colicins (10). FtsH must be proteolytically active for PyoG to translocate into the cytoplasm. Inner membrane transport of PyoG could be coupled with FtsH-dependent proteolytic processing that releases the C domain into the bacterial cytoplasm, as previously demonstrated for nuclease colicins(14, 18). **C** – Like nuclease bacteriocins, nuclease T6SS effectors contain the IMT domain (C – cytotoxic domain, IMT – inner membrane translocation domain, N – other domains and motifs in the N- terminus). The needle and needle tail proteins can deliver these effectors to the *cis* side of the peptidoglycan layer (25). How the nuclease effectors cross the inner membrane and reach the cytoplasm is unknown. It is possible that, like in the case of nuclease PyoG, this translocation step depends on the IMT domain.

We showed that some T6SS effectors, such as the *E. coli* nuclease Usp effector (23), share structural similarities with nuclease bacteriocins. Both groups of toxins can have an HNH motif in the nuclease domain, and the conserved IMT domain upstream of the nuclease domain (Figure 1). Therefore, nuclease bacteriocins and T6SS effectors may share similarities in their inner membrane translocation mechanism. T6SS effectors are loaded to the T6SS needle or needle tail proteins, which penetrate the outer membrane, reach the *cis* side of the peptidoglycan layer (24), and deliver the effector to the bacterial periplasm (25). While the T6SS effectors are directly delivered to the *cis* side of the peptidoglycan layer, bacteriocins may require the pulling force of TonB1 to reach this cell envelope compartment, in a mechanism similar to the one exerted by TolA on TolB and Pal (26). How T6SS effectors further translocate the inner membrane is unknown. It is possible that, like in the case of PyoG, the IMT domain of T6SS effectors is required for inner membrane translocation (Figure 4 – C). Potentially, this translocation also requires FtsH, but this has yet to be tested experimentally.

The conserved IMT domain has previously been linked to bacteriocin killing activity (6), but until now its role in bacteriocin import has been unknown. In this study, we demonstrate that the IMT domain is specifically required for inner membrane translocation of nuclease bacteriocins. We also found this domain in T6SS nuclease effectors, which offers a link between nuclease effector import in contact dependent and independent competition systems. While these systems use different mechanisms to deliver cytoplasmic effectors into the bacterial periplasm, the transport of the effector across the inner membrane may be conserved.

## Materials and Methods

### Sequence analysis of IMT domain containing proteins

IMT domain (PF06958.7) containing proteins were identified in uniref100 using HMMer. Domains were identified using Pfam 27 with e-value < 1x10^-10^. Taxonomy of species with IMT domain was determined using NCBI Commontree and visualised using iTOL (27).

### Bacterial strains, media and growth conditions

All strains (Table 1) were cultured in LB (10 g/L tryptone, 10 g/L NaCl, 5 g/L yeast extract, pH 7.2) or M9 media (8.6 mM NaCl, 18.7 mM NH4CL, 42.3 mM Na2HPO4, 22.0 mM KH2PO4, 0.4% w/v glucose, 2 mM MgSO4, 0.1 mM CaCl2) at 37 °C with shaking (140 rpm). *P. aeruginosa ΔftsH* was grown on salt free LB media. *ΔtonB1* mutant of *P. aeruginosa*, and the parent strain PAO6609, were grown in LB media supplemented with 100 µM FeCl3. *P. aeruginosa Δhur* mutant was grown in the presence of 10 µg/mL tetracycline. 100 µg/mL of carbenicillin and 25 µg/mL of triclosan was used for selecting plasmid transformants of *P. aeruginosa*.

**Table 1.** List of strains used in this study.

| Species | Strain | Characteristics | Source |
| --- | --- | --- | --- |
| *Escherichia coli* | BL21(DE3) | Expression of His-tagged bacteriocins and TonB1 | New England Biolabs |
| | NEB5α | Plasmid propagation | New England Biolabs |
|  | S17-1 | Conjugation with *P. aeruginosa* | American Type Culture Collection |
| *Pseudomonas aeruginosa* | PAO1 | wild type | Washington Library (29) |
|  | PW3356 | PA1302 (*hur*) |  |
| | PAO1 Δ*ftsH* | *ftsH* deletion mutant | (30) |
| | pFtsH | Δ*ftsH* mutant of PAO1 complemented with pREN144 | This study. |
| | pFtsH H416Y | Δ*ftsH* mutant of PAO1 complemented with pREN145 | |
|  | PAO6609 | *met-9011 amiE200 strA pvd-9* | (31) |
|  | PAO6609<br>*ΔtonB1* | *tonB1* transposon mutant |  |
|  | pTonB1 | *ΔtonB1* mutant of<br>PAO6609 complemented<br>with *ptonB1* | This study. |

### Plasmids

pET21a(+) was used as a backbone for the expression of ColD and PyoG constructs in *E. coli* BL21(DE3). pMMB190 was used for complementation and gene expression in *P. aeruginosa*. All plasmids are listed in Table 2. Full length PyoG, with the addition of a G4S linker and a cysteine in the C-terminus in an operon with the ImG immunity protein, was synthesized by Genewiz, and cloned into pET21a(+). pNGH262 (5) was used as the backbone for the construction of plasmids carrying PyoG constructs for the fluorescent labelling of *P. aeruginosa*.

**Table 2.** List of plasmids used in this study.

| Plasmid | Insert | Vector | Description | Source |
| --- | --- | --- | --- | --- |
| pET21a(+) |  |  | pBR322 origin,<br>Hist tag, Amp<sup>r</sup> | NEB |
| pMMB190 | | | Broad-host-<br>range cloning<br>vector,<br>Amp<sup>r</sup>,<br>pMMB66EH,<br>tac promoter,<br>LacZα | (32) |
| pNGH262 | PyoG, ImG-His<sub>6</sub> | pET21a(+) | His<sub>6</sub>-Im-PyoG<br>cloned at *Nde*I<br>and *Hind*III sites | (5) |
| pNGH263 | ImG-His<sub>6</sub> | pACYCDuet1 | PyoG immunity<br>protein cloned<br>at *Nde*I and<br>*Xho*I | (5) |
| pG-G4S-Cys | PyoG-G4SC,<br>ImG-His<sub>6</sub> | pET21a(+) | PyoG with a<br>G4S linker and<br>cysteine in the<br>C-terminus | This<br>study |
| pG1-485-Cys | PyoG<sup>1-</sup><br><sup>485</sup>Cys-His<sub>6</sub> | pET21a(+) | PyoG<sup>1-485</sup>-Cys-His<sub>6</sub> cloned at<br>*NdeI* and<br>*HindIII* sites | This<br>study |
| pG1-255-Cys | PyoG<sup>1-</sup><br><sup>255</sup>Cys-His<sub>6</sub> | pET21a(+) | PyoG<sup>1-255</sup>-Cys-His<sub>6</sub> cloned at<br>*NdeI* and<br>*HindIII* sites | This<br>study |
| pREN144 | His6-TEV-FtsH | pMMB190 | *ftsH* from PAO1<br>with an N-terminal His6<br>and TEV site | This<br>study |
| pREN145 | His6-TEV-FtsH, point<br>mutation<br>H416Y | pMMB190 | *ftsH* H416Y<br>from PAO1 with<br>an N-terminal<br>His6 and TEV<br>site | This<br>study |
| pTonB1 | *tonB1* | pMMB190 | *tonB1* from<br>PAO1 genome | This<br>study |

pG1-485-Cys was constructed by PCR, amplifying the first 485 residues of PyoG and by adding a cysteine at the C-terminus of the construct by PCR mutagenesis (5). pG1-255-Cys was constructed by introducing a cysteine and an XhoI site after residue A255 in pNGH262 and cutting out the region between the two XhoI sites. Genes used for *P. aeruginosa* complementation (*ftsH*, *ftsH* H416Y and *tonB1*) were synthetised by Genewiz and restriction cloned into pMMB190 using the EcoRI and HindIII sites. Plasmid DNA was isolated from NEB5α cultures grown at 37 °C in LB with the appropriate antibiotic, using Monarch Plasmid Miniprep kit (NEB). Sequencing was through the company Genewiz.

### Conjugation of *P. aeruginosa*

Chemically competent *E. coli* S17-1 competent cells were prepared by CaCl2 treatment. 50 mL of over-night culture, grown at 37°C in LB, was pelleted on 5000 x g for 10 min. Cells were resuspended in 50 mL ice cold 0.1 M CaCl2 and kept on ice for 30 min. Cells were then pelleted again and resuspended in 1 mL 0.1 M CaCl2 and incubated on ice for 5 more min before transformation. Competent cells were transformed by heat shock. 50 µL of competent cells were mixed with 5 µL of DNA. Cells were incubated on ice for 30 min, shocked for 30 s on 42 °C, and then incubated on ice for another 10 min. Cells were then plated on LB agar with antibiotics. *P. aeruginosa* PAO1 was transformed via conjugation with *E. coli* S17-1 carrying a plasmid of interest. PAO1 was grown overnight at 42 °C, and S17-1 at 37 °C, with shaking, in LB media. 2 mL of each culture was pelleted on 5000 x g, 10 min. Each strain was then resuspended in 100 µL of LB media and the two strains were mixed together. The entire mix was then spotted on top of an antibiotic free LB agar plate. The plate was incubated on 37 °C for 4 h. The lawn was then scraped from the plate and resuspended in 1X phosphate buffer saline (PBS) pH 7. The suspension was serially diluted in PBS. 100 µL of each dilution was plated on LB agar with 25 µg/mL triclosan, for selecting against S17-1, and 100 µg/mL carbenicillin, for selecting against untransformed PAO1.

### Expression and purification of bacteriocin constructs

PyoG was expressed in complex with the His-tagged immunity protein, ImG. Two copies of ImG were used: one in an operon with PyoG (pNGH262), and one on a separate plasmid (pNGH263), as described previously (5). PyoG^1-485^ and PyoG^1-255^ were expressed with an N-terminal His-tag. *E. coli* BL21(DE3) was used for heterologous pyocin expression, as described previously (5).

### Circular dichroism (CD) spectroscopy

Proteins were dialysed into 10 mM potassium phosphate pH 8, 20 mM NaF, and were diluted to 0.1 mg/ml. CD spectra were obtained using a Jasco J-815 Spectropolarimeter over a wavelength range of 260-190 nm, a digital integration time of 1 second and a 1 nm bandwidth. CD data in millidegrees were converted to mean residue ellipticity by dividing by molar concentration and number of peptide bonds.

### Differential scanning calorimetry (DSC)

The melting temperature of proteins, as an indication of their integrity, was determined by DSC performed on Malvern VP Capillary DSC by Dr. David Staunton, Molecular Biophysics Suite, Department of Biochemistry, University of Oxford. Pyocin constructs were tested at 20 µM in 10 mM potassium phosphate pH 8, 20 mM NaF.

### Conjugation of maleimide fluorophores to proteins

Pyocins were fluorescently labelled using Alexa Fluor 488 C5 maleimide (AF488) fluorophore that was linked to proteins via an engineered C-terminal cysteine, as previously described (5).

### Labelling of live *P. aeruginosa* cells with fluorescent pyocins

Fluorescent PyoG constructs conjugated to AF488 were used to label *P. aeruginosa*. Bacteria were grown overnight in M9 medium at 37 °C with shaking. 1 mL of this overnight culture was pelleted and resuspended in 10 mL M9 medium and grown until an OD600 of 0.5. All pelleting steps were performed at 7000 x g for 3 min. 1 mL of cells was washed in PBS pH 7, and labelled with 2 µM pyocin for 30 min at room temperature. The unbound pyocin was removed by three washes in PBS. For the trypsin protection assay, after labelling, cells were exposed to 0.5 mg/mL trypsin for 1 h at 30 °C, in PBS with 35 µg/mL chloramphenicol. After a final washing step, bacteria were resuspended in 30 µL of PBS. 3 µL of cells were then loaded onto agarose pads, prepared using Geneframes (Thermo Scientific). 80 µL of 1% (w/v) agarose in PBS was pipetted into the Geneframe (17×28 mm). The surface was flattened with a cover slip and excess agar removed. Once the agar solidified the cover slip was removed, the bacterial suspension added and a new coverslip attached to the adhesive side of the Geneframe.

### Labelling of *P. aeruginosa* spheroplasts with fluorescent pyocins

Bacteria were grown overnight in M9 medium at 37 °C with shaking. 1 mL of this overnight culture was pelleted and resuspended in 10 mL M9 medium and grown until an OD600 of 0.5. All pelleting steps were performed at 3000 x g for 10 min. 1 mL of cells was pelleted and resuspended in PBS pH 7, with the addition of 0.5 M sucrose, 20 mM EDTA and 1.5 mg/mL lysozyme, and incubated for 45 min on room temperature. Cells were then washed into PBS, 0.5 M sucrose, and mixed with 2 µM fluorescent pyocin. After 30 min incubation on room temperature, unbound pyocin was removed by three washes in PBS, 0.5 M sucrose. For the trypsin protection assay, after labelling, cells were exposed to 0.5 mg/mL trypsin for 1 h at 30 °C, in PBS, 0.5 sucrose. After a final washing step, bacteria were resuspended in 30 µL of PBS, 0.5 M sucrose and loaded onto agar pads as described above. Agar was supplemented with 0.5 sucrose to prevent the bursting of the spheroplasts.

### Image collection and data analysis

All images were collected on an Oxford Nanoimager S microscope. In case of PyoG constructs, images were collected at 100 ms exposure and 20 % 488 nm laser power. For every image, 20 frames were collected and merged using the command “Zproject” in Image J. Average fluorescence was measured for a total of 50 cells per condition per repeat, and was corrected by subtracting the average background fluorescence. All experiments were conducted in 3 biological repeats. Fluorescence intensity of experimental groups (groups exposed to fluorescent pyocin) was compared to the unlabelled control by the Kruskal-Wallis test, using Dunn’s test as the post hoc procedure (confidence level 0.001). The analysis was performed by GraphPad Prism version 6.04 for Windows, GraphPad Software, La Jolla California USA, www.graphpad.com.

### Plate killing assays

*P. aeruginosa* was grown in LB at 37 °C to an OD600 of 0.6. Bacterial lawns were prepared by addition of 250 µl of culture to 5 ml of molten soft LB-agar (0.75% (w/v) agar in LB), and were poured over LB-agar plates. Once set and dry, 3 µl of 3-fold serially diluted pyocins, ranging from 10 µM to ∼57 pM, was spotted on top of the lawn. Lawns were grown overnight at 37 °C and cytotoxicity was determined by observation of clearance zones.

## Acknowledgments

The authors thank Dr Nick Housden, Dr Joanna Szczepaniak, Dr Ruth Cohen- Khait, Dr Nathalie T Reichmann and Dr Melissa Webby for useful discussions. We also thank Dr David Staunton of the Molecular Biophysics Suite (Department of Biochemistry) for help with all biophysical experiments. I.A. was funded by the Wellcome Trust Infection, Immunology and Translational Medicine Doctoral Training Centre. This work was supported by a Wellcome Trust Collaborative award (201505/Z/16/Z) funded to C.K.

---

**Supplementary Figure 1.**
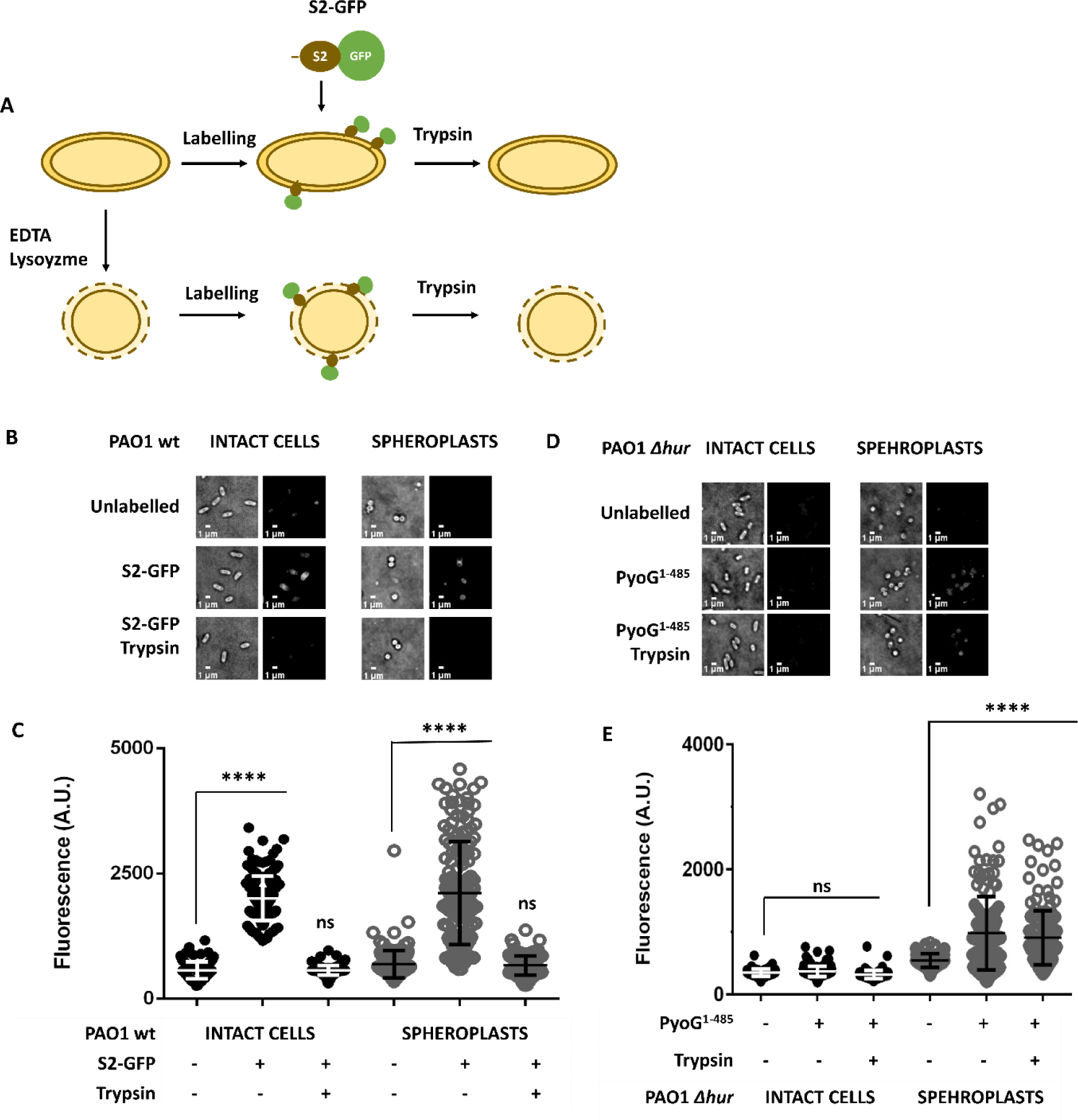
S2-GFP is not trypsin protected in *P. aeruginosa* PAO1 intact cells and spheroplasts. **A** – S2-GFP is comprised of the S2 N-terminal domain (S2^1-209^), translationally fused to GFP. GFP acts as a plug that blocks translocation of S2 across the cell envelope (10). This construct is used to test if surface exposed pyocin gets degraded by trypsin in the fluorescence microscopy experiment described in Figure 1. **B, C** - Intact cells, or spheroplasts generated with lysozyme/EDTA treatment, are exposed to 2 µM S2-GFP. The S2-GFP fluorescent signal is removed by trypsin, in both intact cells and spheroplasts. This indicates that untranslocated pyocin is removed by trypsin under tested experimental conditions. **D, E** - Hur, the outer membrane receptor of PyoG, is not required for PyoG import into spheroplasts. Therefore, the outer membrane translocation step is bypassed in spheroplasts, which enables specific detection of inner membrane transport. Intact cells of PAO1 *Δhur* are not labelled with the pyocin, since Hur is required for binding of PyoG to the surface of *P. aeruginosa* cells (5). Generation of spheroplast with lysozyme/EDTA treatment is sufficient to allow *Δhur* labelling with fluorescent PyoG. **B, D** - Representative micrographs before and after trypsin treatment are shown. All snapshots were adjusted to the same intensity scale. **C, E** - Average fluorescence intensities were measured for 150 cells per condition. Mean of three biological replicates with standard deviations are shown. Fluorescence intensities for labelled and trypsin treated groups in each condition are compared to the unlabelled control. **** represents a P value below 0.0001 in the Kruskal-Wallis Test, and ns represents no significant difference or lack of fluorescent labelling.

**Supplementary Figure 2.**
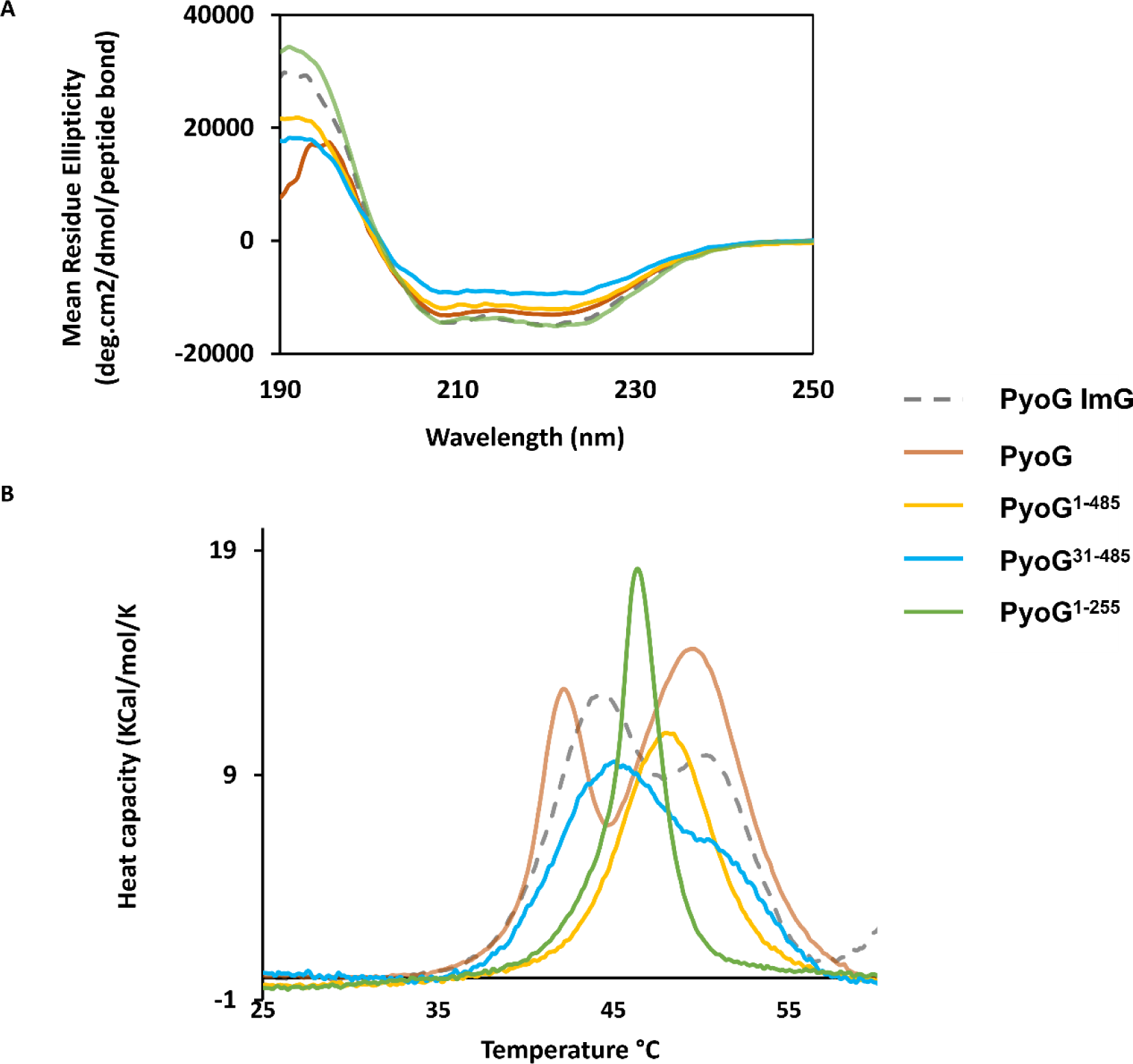
Quality control of PyoG construct used for fluorescent labelling of *P. aeruginosa*. **A** - CD spectra of 0.1 mg/ml PyoG-ImG (dashed line), PyoG (orange), PyoG^1-485^ (yellow), PyoG^31-485^ (blue), and PyoG^1-255^ (green) at room-temperature in 20 mM NaF and 10 mM KPO4 buffer (pH 7). PyoG has a G4S linker and a cysteine at the C terminus. All the other constructs have a cysteine directly added to the C-terminus. An average of 9 measurements are shown. **B** - DSC curve of 20 µM PyoG constructs in 50 mM Tris-HCl pH 7, 150 mM NaCl, in the 25-60 °C temperature range. The melting temperature of PyoG is 42.14 ± 0 and 49.56 ± 0.02 °C, PyoG^1-485^ is 48.04 ± 0.14 °C, and PyoG^1-255^ is 46.22 ± 0.02 °C. All constructs have a similar melting curve to the wt PyoG-ImG, indicating that they are folded at room temperature.

**Supplementary Figure 3.**
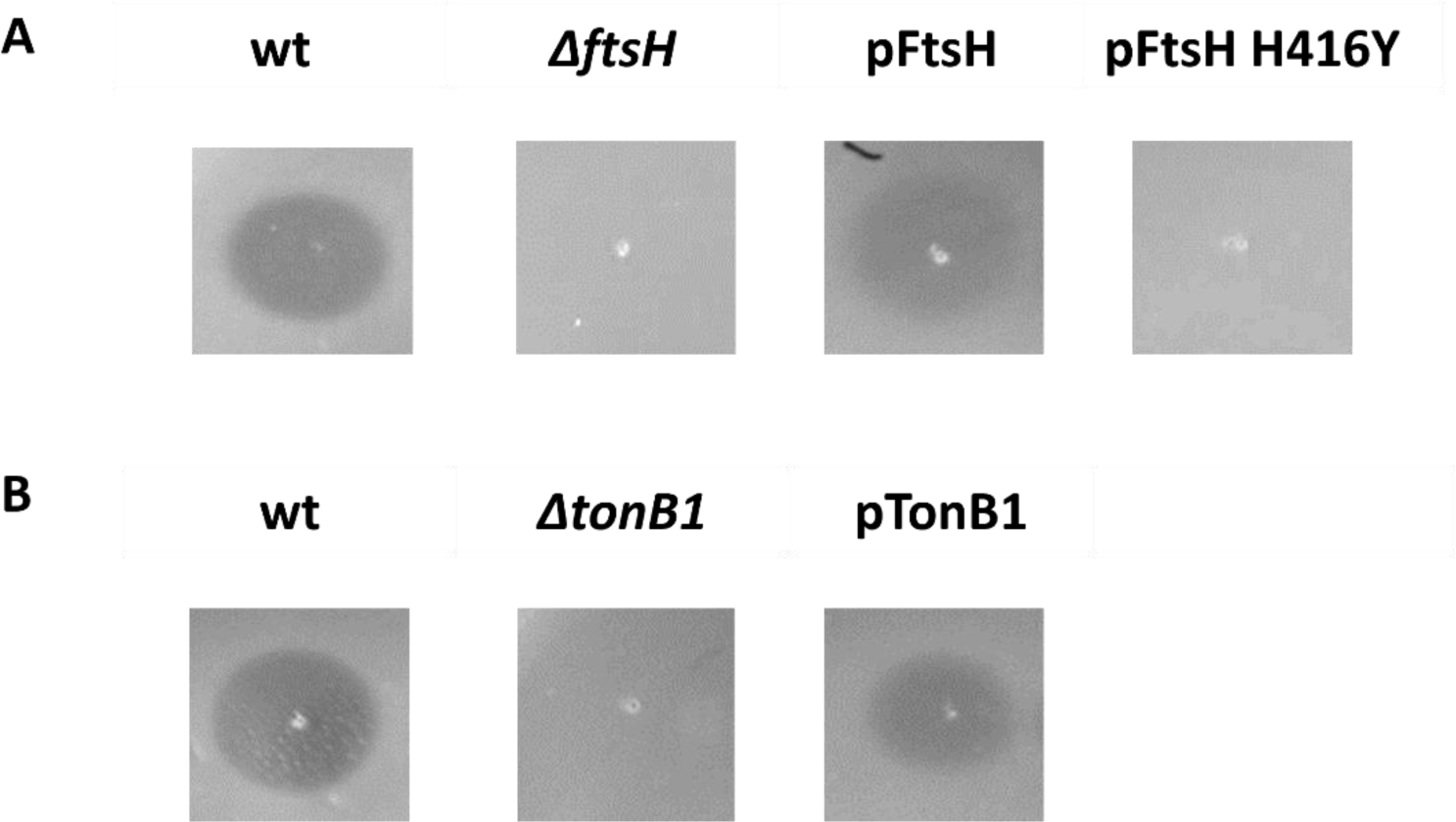
Plate killing assay of *P. aeruginosa* complemented strains. **A** - PAO1 *ΔftsH* complemented with FtsH expressed from a plasmid (pFtsH), or a protease inactivated version of FtsH (pFtsH H416Y). Only complementation with wt FtsH restored PyoG sensitivity. **B** - *P. aeruginosa* PAO6699 *ΔtonB1* was complemented with TonB1 expressed from a plasmid (pTonB1). Pyocin sensitivity is restored in this strain.

**Supplementary Figure 4.**
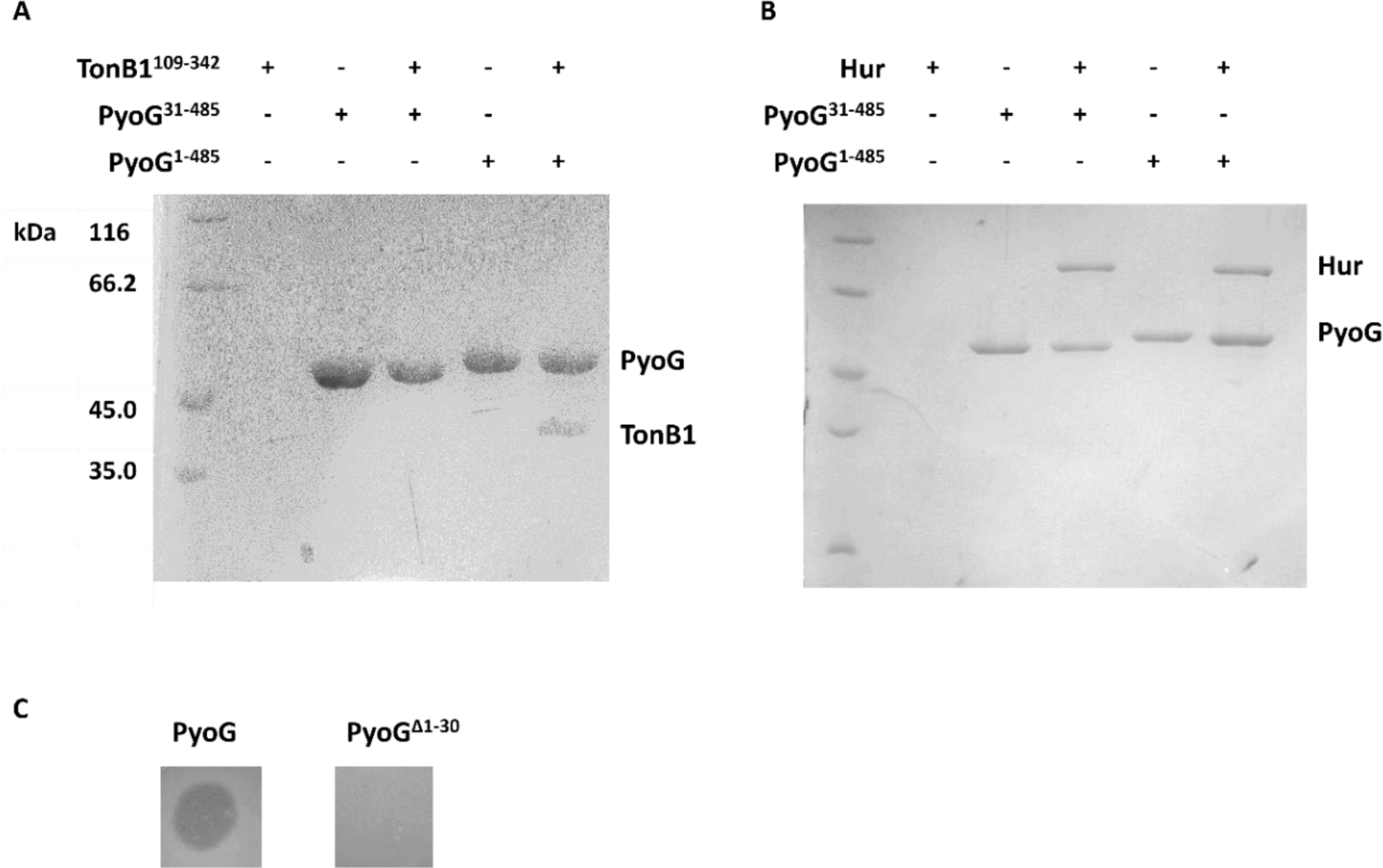
Deletion of the first 30 residues of PyoG disrupts TonB1 binding and the pyocin killing activity. **A, B** - Pulldowns with the PyoG^31-485^ construct. This construct was used for fluorescent labelling of *P. aeruginosa* (Figure 3 **- G, H**). It has a cysteine and a His6-tag in the C-terminus and was used as bait protein in pull-down assays. Periplasmic TonB1 (**A**) and Hur (**B**) were purified as described previously (5). Both proteins had no purification tag and were used as prey proteins. All proteins were mixed to a final concentration of 10 µM in binding buffer (50 mM Tris-HCl pH 7.8, 250 mM NaCl; 1% β-OG was added if Hur was used) and bound to nickel beads on room temperature. Unbound protein was washed in the same buffer and bound proteins were eluted in the presence of 250 mM imidazole. Eluate was analysed on 12% SDS-PAGE gels. Proteins that were added to beads are indicated above each lane. Positions of proteins are labelled on the right side of each gel. PyoG^31-485^ does not bind TonB1 in the pull-down assay (**A**), but it binds to Hur (**B**). This indicates that the first 30 residues of PyoG contain the TonB1 binding box, and that this region is not essential for receptor binding. **C** – The killing activity of PyoG which lacks the first 30 residues (PyoG^Δ1- 30^). Unlike PyoG^31-485^, PyoG^Δ1-30^ contains the cytotoxic domain, and its activity can be tested in a plate killing assay. 3 µL drops of 10 µM wt and Δ1-30 PyoG were spotted on PAO1 lawns. The deletion of the first 30 residues of PyoG disrupted its killing activity against PAO1, which indicates that this region of the pyocin is essential for its import into bacterial cells.

